# Convergent evolution of p38/MAPK activation in hormone resistant prostate cancer mediates pro-survival, immune evasive, and metastatic phenotypes

**DOI:** 10.1101/2020.04.22.050385

**Authors:** Kathryn E. Ware, Santosh Gupta, Jared Eng, Gabor Kemeny, Bhairavy J. Puviindran, Wen-Chi Foo, Lorin A. Crawford, R. Garland Almquist, Daniella Runyambo, Beatrice C. Thomas, Maya U. Sheth, Anika Agarwal, Mariaelena Pierobon, Emanuel F. Petricoin, David L. Corcoran, Jennifer Freedman, Steven R. Patierno, Tian Zhang, Simon Gregory, Zoi Sychev, Justin M. Drake, Andrew J. Armstrong, Jason A. Somarelli

## Abstract

Adaptation of cancer cells to targeted therapy follows ecological paradigms observed in natural populations that encounter resource depletion and changing environments, including activation of pro-survival mechanisms, migration to new locations, and escape of predation. We identified the p38 MAPK pathway as a common molecular driver of these three responses during the adaptation to hormone therapy resistance in prostate cancer. The p38 pathway is activated in therapy-resistant cells and mechanistically drives these three convergent responses through sustained AR activity, enhanced invasion and metastasis, and immune evasion. Targeting p38 signaling may represent a new therapeutic strategy to treat men with metastatic, hormone therapy-resistant prostate cancer.

## Introduction

Populations of individuals within an ecological niche must acquire the necessary resources to survive and propagate their genetic material to the next generation. Individuals have adapted a wide range of strategies to ensure resource acquisition in an ever-changing environment. Some of these strategies include dormancy (Varpe, 2017), hibernation (KIlduff, 2004), migration, and avoidance of predation (Skov et al., 2010).

Cancer cells adapt similar strategies within the context of the ecological niche of the body to cope with variations in resource availability that promotes their survival. A major resource for cancer cells is the oncogene signaling pathway to which they are addicted. For prostate cancer cells, this critical lineage oncogene is most often the androgen receptor (AR) (Mills, 2014). AR activity is pharmacologically targeted through blocking the ligand binding domain or inhibition of androgenic ligand biosynthesis. Enzalutamide, a 2^nd^ generation inhibitor of the androgen receptor, and abiraterone acetate, an androgen synthesis inhibitor, each delay progression and improve the survival of men with both early and late castration-resistant prostate cancer (Beer et al., 2014; de Bono et al., 2011; Penson et al., 2016; Ryan et al., 2013; Scher et al., 2012). These potent hormonal therapies significantly prolong the overall survival of men with metastatic, castration-resistant prostate cancer; however, acquired resistance to these drugs over a median of one to two years is inevitable.

Upon disease progression with enzalutamide or abiraterone treatment, most tumors remain AR dependent and have a rise in serum levels of prostate-specific antigen (PSA) (Bluemn et al., 2017; Bryce et al., 2017). Multiple mechanisms of AR signal restoration have been identified that directly impact the AR gene, including AR amplification, AR mutations, genomic structural rearrangements (Li et al., 2011b; Li et al., 2012; Liu et al., 2013; Ware et al., 2014), and alternative splicing events (Liu et al., 2014). Additionally, alternative mechanisms can promote the AR transcriptome or promote AR activity through substitute methods of AR activation (Arora et al., 2013; Ware et al., 2014) to generate a pro-survival response (Chen et al., 2004; Viswanathan et al., 2018).

In addition to AR activation, complementary pro-survival responses focus on metabolic plasticity, such as dormancy and hibernation. Cancer cells adapt their energetic needs to accompany survival and fitness in hostile environments (Lehuede et al., 2016), and disseminated tumor cells can be found in many prostate cancer patients prior to any clinical symptoms (Ruppender et al., 2013; van der Toom et al., 2016). Furthermore, organisms and cancer cells alike must avoid predation to ensure their survival in any environment. Prostate cancer cells avoid the predation of the immune system in a number of ways, including 1) secretion of immunosuppressive molecules, such as TGF-β (Yang et al., 2010; Yoshimura and Muto, 2011) and soluble WNT ligands (Robinson et al., 2015), and 2) expression of cell surface or cellular immune checkpoint molecules (Antonarakis et al., 2020; Gao et al., 2017; Graff et al., 2016). For example, TGF-β has been identified as a potent immunosuppressive ligand, which can be regulated through Snail to mediate downregulation of HLA-I and promote immune escape (Chen et al., 2015). Similarly, the immunosuppressive ligand PD-L1 is upregulated in response to enzalutamide-resistant progression, both in tumor cells and in circulating immune subsets (Bishop et al., 2015; Graff et al., 2016). Additionally, we have shown that PD-L1 expression is more prevalent on circulating tumor cells from metastatic prostate cancer patients who are progressing on abiraterone acetate or enzalutamide treatment (Zhang et al., 2018).

In the present study we sought to understand the adaptations to hormone therapy resistance in prostate cancer. Using an integrated genomics approach we observed that enzalutamide-induced AR signaling blockade induces convergent phenotypic evolution on three ecological responses: 1) altered resource acquisition to promote cellular persistence and survival through oncogene re-activation, 2) upregulation of migratory/invasive factors, and 3) avoidance of predation by immune evasion and immune suppression (Fig. 1). All three of these phenotypes converge across different model systems on the p38/MAPK stress response pathway, which is highly activated in human prostate cancer metastases and can be therapeutically leveraged to simultaneously target and limit these pro-survival responses to overcome enzalutamide resistant growth and survival.

**Figure 1.**
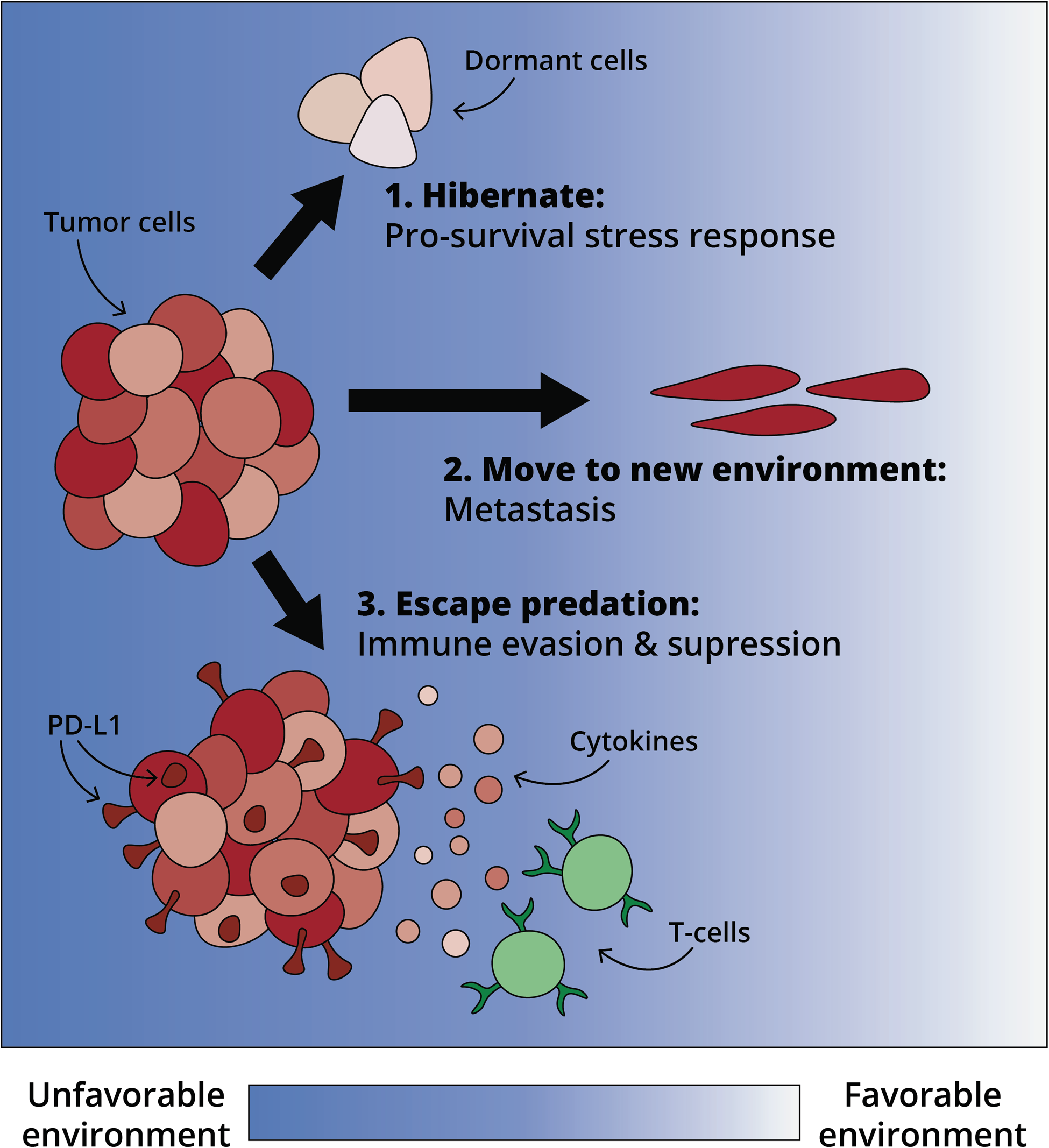
Model of ecological responses to loss of AR signaling. Adaptation to enzalutamide centers around three phenotypes: 1) Pro-survival stress response; 2) Immune evasion; and 3) Metastasis.

## Results

### Enzalutamide resistant cells exhibit diverse genomic adaptations

Enzalutamide is a potent inhibitor of AR activity (Tran et al., 2009) that initially induces a response in most men with metastatic castration resistant prostate cancer; however, progression on enzalutamide typically develops within 1-2 years (Beer et al., 2014). A deeper understanding of the mechanisms underlying the evolution of enzalutamide resistance is needed to target these resistance mechanisms. To identify common molecular mechanisms of enzalutamide resistance we developed a panel of four enzalutamide-resistant (enzaR) cell lines, LNCaP-enzaR, CS2-enzaR, LN95-enzaR, and MDA-PCa-2b-enzaR, by chronic, long-term exposure to increasing doses of enzalutamide. For LNCaP and MDA-PCa-2b cells, the cells were initially exposed to 1 μM enzalutamide and allowed to grow to confluence. The dose of enzalutamide was doubled at each subsequent passage until cells were capable of sustained growth in the presence of 50 μM enzalutamide (Fig. 2A), which is above the concentration observed in men with metastatic, castration resistant prostate cancer (Scher et al., 2010). In parallel with development of these enzalutamide-resistant models, we also created enzalutamide-resistant populations of LNCaP sublines, CS2 and LN95. Both CS2 and LN95 (Hu et al., 2012) cells were first adapted to androgen deprivation therapy (ADT) by passaging in media supplemented with an increasing ratio of androgen-depleted media. Following evolution of ADT resistance, cells were then adapted to enzalutamide with increasing doses as described above (Fig. 2A). All enzaR cell lines are characterized by a significant increase in cell growth and a decrease in apoptosis in response to enzalutamide treatment (Fig. 2B).

**Figure 2.**
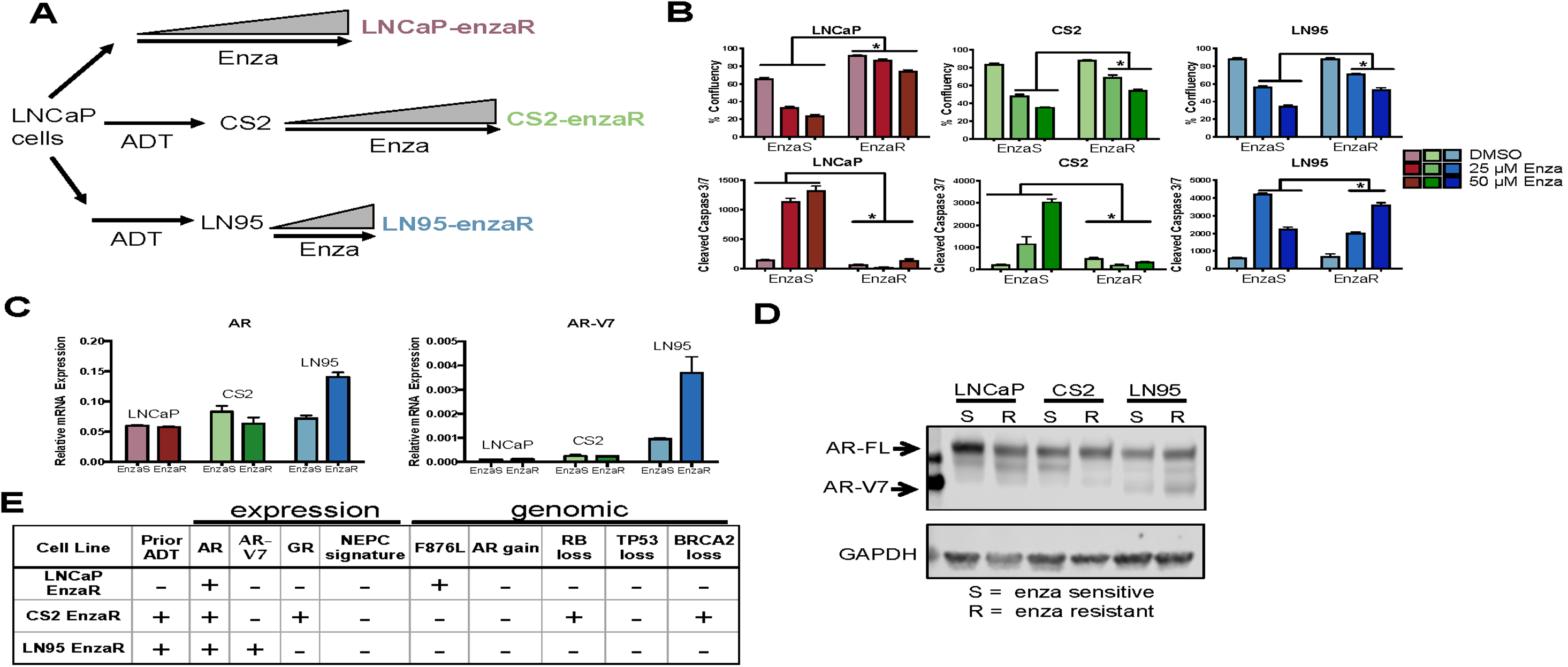
Enzalutamide resistant cell lines retain AR expression and do not undergo a NEPC/EMT/Stem transition. **A.** Development of enzalutamide resistant cell lines. **B.** Control (ctrl) and enzaR cell lines were treated for 10 days with vehicle (DMSO) or two doses of enzalutamide (25μM or 50μM), and cell growth and apoptosis were measured. *= p<0.05; student’s t-test. Error bars represent the standard error of the mean. **C-D.** AR-FL and AR-V7 mRNA (**C**) and protein expression (**D**) measured by qRT-PCR and western blot, respectively, *p<0.05; student’s t-test. Mean cell values are plotted along with standard error of the mean. **E.** Summary table characterizing the diversity of known resistance mechanisms in enzalutamide resistant cell lines. ADT, androgen deprivation therapy; Enza, enzalutamide; EnzaR, enzalutamide resistant.

Analysis of these enzaR cells revealed several clinically-relevant genotypes and phenotypes. All cell lines had persistent AR expression (Fig. 2C-D) and did not have notable morphologic changes or consistent neuroendocrine differentiation (**Supplementary Fig. S1A**). The enzalutamide-resistant LNCaP model acquires the F876L mutation in AR that converts enzalutamide into a partial agonist (Balbas et al., 2013; Korpal et al., 2013) (**Supplemental Fig. S2**), but lack additional novel AR mutations. The enzaR LNCaP and CS2 cells express full-length AR, but do not produce the AR splice variant, AR-V7, which is a known enzalutamide-resistance driver (Antonarakis et al., 2014) (Fig. 2C-E). On the other hand, enza-R LN95 cells upregulate both full length AR and AR-V7 (Fig. 2C-E). Enza-R CS2 cells display upregulation of the glucocorticoid receptor mRNA and genomic loss of *RB1* and *BRCA2*, which are known drivers of resistant and aggressive prostate cancer (Arora et al., 2013; Chakraborty et al., 2019). None of these lines have upregulation of biomarkers of neuroendocrine lineage plasticity as measured by co-*RB1* and *TP53* genomic loss or transcriptional upregulation of *SOX2*, and none of these lines expresses markers of neuroendocrine lineage plasticity (Mu et al., 2017) at the mRNA level (**Supplemental Fig. S1B**). We also did not observe changes in the stemness markers *CD133, NANOG*, or *OCT4* in the enzaR cell lines (Fig. 2E, **Supplemental Fig. S3**). Thus, this panel of cell lines parallels the heterogeneous spectrum of AR-positive prostate cancer phenotypes most commonly observed in the clinic upon progression on enzalutamide.

To further understand the evolution of enzalutamide resistance in these heterogeneous models of prostate cancer, we analyzed the genome, transcriptome and phospho-proteome of these paired cell lines prior to enzalutamide exposure and following adaptation to enzalutamide resistance. Genetically, the enzaR models were all remarkably unique. While enzaR CS2 and LN95 cells share just one copy number alteration – a gain in chromosome 20p13-p11.1 – this gain is not present in enzaR LNCaP cells (**Supplemental Fig. S4**). Likewise, enzaR models acquired relatively few single nucleotide variants in the exome between paired parental and enzaR cell lines, and of these alterations, none were shared across all enzaR cells (**Supplemental Fig. S5A-B**). Together, these data suggest that the enzaR lines do not harbor shared genetic alterations that contribute to their adaptation to enzalutamide.

### Enzalutamide resistance converges on MAPK signaling and stress response pathways

Given the diversity of genetic lesions in the enzaR models, we next sought to determine the molecular drivers underlying the evolution of enzalutamide resistance. Across each parental and resistant model system, we performed whole transcriptome RNA Sequencing to identify consistent changes in RNA expression and pathways associated with resistance. Similar to our DNA-level analyses, we observed relatively few gene expression changes common to all cell line models (4-5% overlap; Fig. 3A). We also performed a reverse phase protein array (RPPA) and evaluated a subset of proteins and phosphoproteins that are implicated in cancer progression (Akbani et al., 2014). The proteomics data revealed phospho-p38, a key signaling node in the cellular stress response, as the top upregulated phospho-protein in all three enzaR cell lines (Fig. 3C). Reanalysis of our RNA-Seq data at the pathway level revealed common transcriptional enrichment of MAPK signaling, DNA repair, and several stress-response pathways (Fig. 3B). Importantly, we observed the same enrichment of stress-response pathways and MAPK signaling in a fourth enzalutamide-resistant cell line, MDA-PCa-2b (Fig. 3D, E). MDA-PCa-2b is an independent androgen responsive and AR positive prostate cancer cell line. These data suggest that AR-positive enzaR cells exhibit unique genetic and transcriptional landscapes at the gene level that converge on p38 signaling and stress response signaling at the pathway level. These pathways are activated as a consequence of adaptation to enzalutamide and not an acute response, as a short-term (5 day) treatment with enzalutamide does not activate this same p38-mediated response pathway (**Supplemental Fig. 6B**). This pathway-level convergence is reminiscent of convergent evolution observed in ecological contexts during resource depletion, including the conserved activation of the p38 MAPK pathway itself (Gatenby et al., 2011; Harrison et al., 2004; Li et al., 2011a).

**Figure 3.**
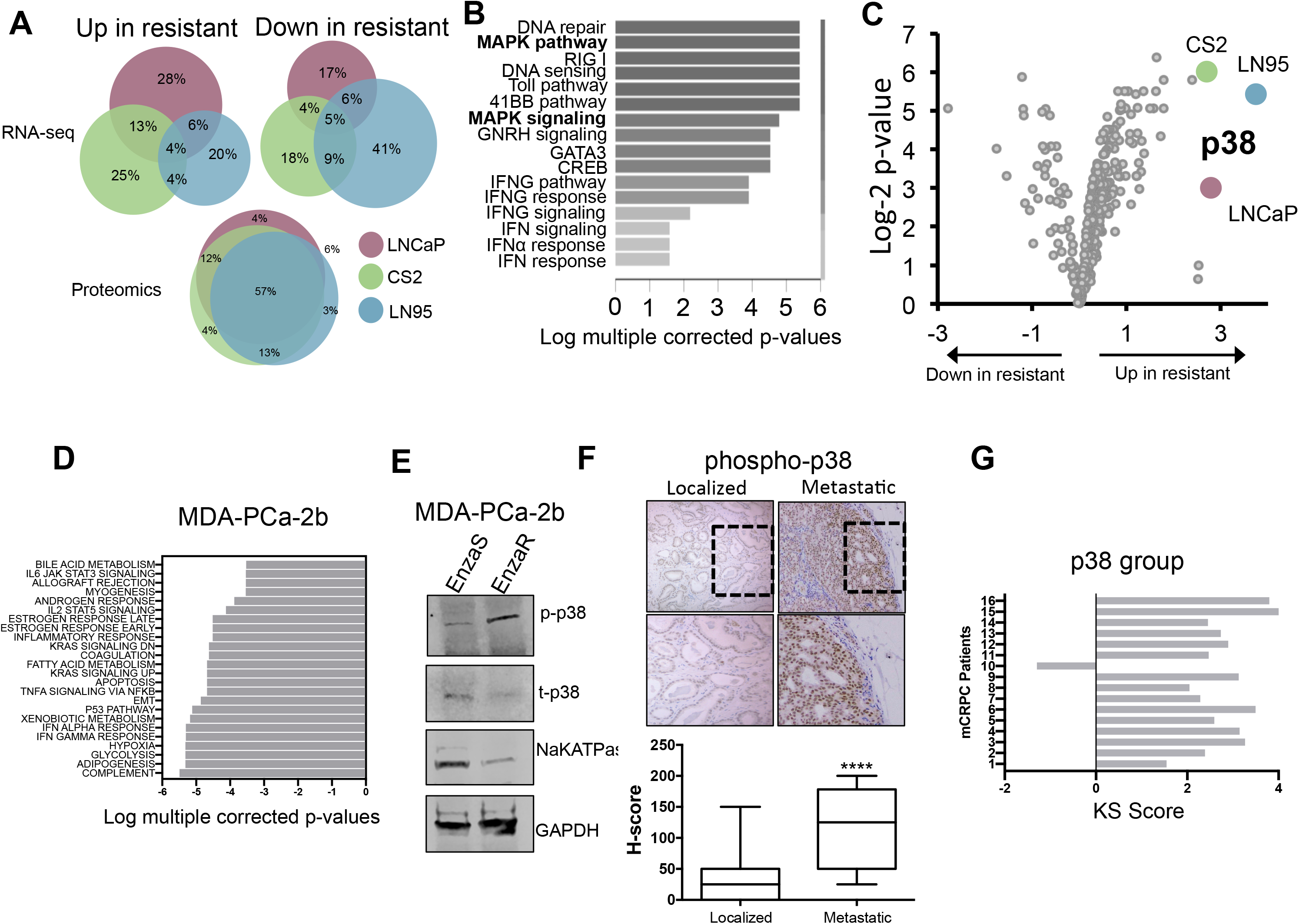
Diverse gene expression during enzalutamide resistance converges on activation of stress response MAPK p38. **A.** Venn diagrams representing common gene expression alterations by RNA-Seq or protein activation by proteomics analysis (top). Enriched pathways in each matched enzaR cell line using gene set enrichment analysis (bottom) from RNA-Seq data. **B.** Combined RNA-Seq analysis from all enzalutamide-resistant lines identified p38/MAPK signaling and stress-related pathways. **C.** Volcano plot representing changes in phosphorylation from reverse phase protein array analysis. **D.** RNA-Seq analysis for pathway enrichment scores in MDA-PCa-2b cells. **E.** Western blot analysis of phosph-p38 in MDA-PCa-2b cells. **F.** Immunohistochemical staining of phospho-p38 is strongly enriched in metastatic tissues compared to localized samples. Representative images show phospho-p38 staining in localized and metastatic prostate cancer tissues. **G.** Kinase substrate enrichment analysis (KSEA) of p-p38 revealed an enrichment of p38 substrates in metastatic CRPC patients using published phosphoproteomic datasets.

Considering the convergent evolutionary behavior of the enzaR cells at the gene expression/signaling levels, we next attempted to better understand if these changes also promoted phenotypic convergence during the acquisition of enzalutamide resistance. To do this, we first explored the phenotypic consequences of p38/MAPK pathway activation. The p38/MAPK pathway is a key regulator of the stress response (Igea and Nebreda, 2015), and a regulator of tumor cell dormancy in many cancer types, including prostate cancer (Decker et al., 2017; Yu-Lee et al., 2018). Consistent with the role of p38 in a dormancy phenotype, enzaR cells exhibited a significant downregulation in the ratios of pERK:p38α, an important indicator of the shift from a proliferative (high ERK) to dormant (high p38α) phenotype (**Supplemental Fig S7A**). Likewise, enzaR cells also upregulated p21, a cyclin dependent kinase inhibitor induced during stress response to regulate cell cycle progression (Sosa et al., 2011) (**Supplemental Fig S7B**). Interestingly, unlike enzalutamide-sensitive cells that induce beta-galactosidase activity, a marker of dormant cells, enzaR cells do not increase beta-galactosidase activity in response to enzalutamide treatment. However, enzaR cells have a higher baseline level of beta-galactosidase activity compared to enzalutamide sensitive cells (**Supplemental Fig S7C**). These data suggest that cells adapt to survival during enzalutamide treatment by regulating cell cycle progression and escaping from treatment induced dormancy programs.

### Activation of p38/MAPK is enriched in metastatic disease and drives cell growth in prostate cancer

Our results indicate that enzaR cells converge on p38 activation, which is functionally associated with dormancy and a pro-survival phenotype. Prior work has suggested activation of the p38 pathway in disseminated prostate and breast cancer cells from patients (Chery et al., 2014; Werden et al., 2016). Based on our preclinical data and these observations, we hypothesized that p38 activation would be associated with aggressive and metastatic disease. To test our hypothesis we compared, by immunohistochemistry, 30 primary tumors and 20 metastatic biopsies from prostate cancer patients (Ware et al., 2016) for phospho-p38 expression. These tumors were all positive for AR expression. Importantly, as compared to localized prostate cancer, metastases show a strong activation of p38 (Fig. 3F).

To validate these findings, we performed kinase substrate enrichment analysis (KSEA) on our published phosphoproteomic dataset (Drake et al., 2016) consisting of 16 metastatic prostate cancer patients. KSEA demonstrated significantly enriched hyperphosphorylation of p38 substrates in metastatic castration-resistant prostate cancer (CRPC) tissues compared to localized hormone-naïve prostate cancer tissue (Fig. 3G, **Supplemental Table 1**). Independent data sets analysis shows p38 activation as the central hub of cell signaling convergence in human prostate cancer metastases, particularly during the evolution of metastasis and hormone therapy resistance.

Next, we tested whether inhibition of p38 signaling impacted cell growth in our models of enzalutamide resistance. In all enzaR cell line models, treatment with a small molecule p38 inhibitor (SB203580) led to a significant decrease in cell proliferation over time (Fig. 4A-C). These data highlight the p38 signaling pathway as important and activated in enzalutamide resistant prostate cancer.

**Figure 4.**
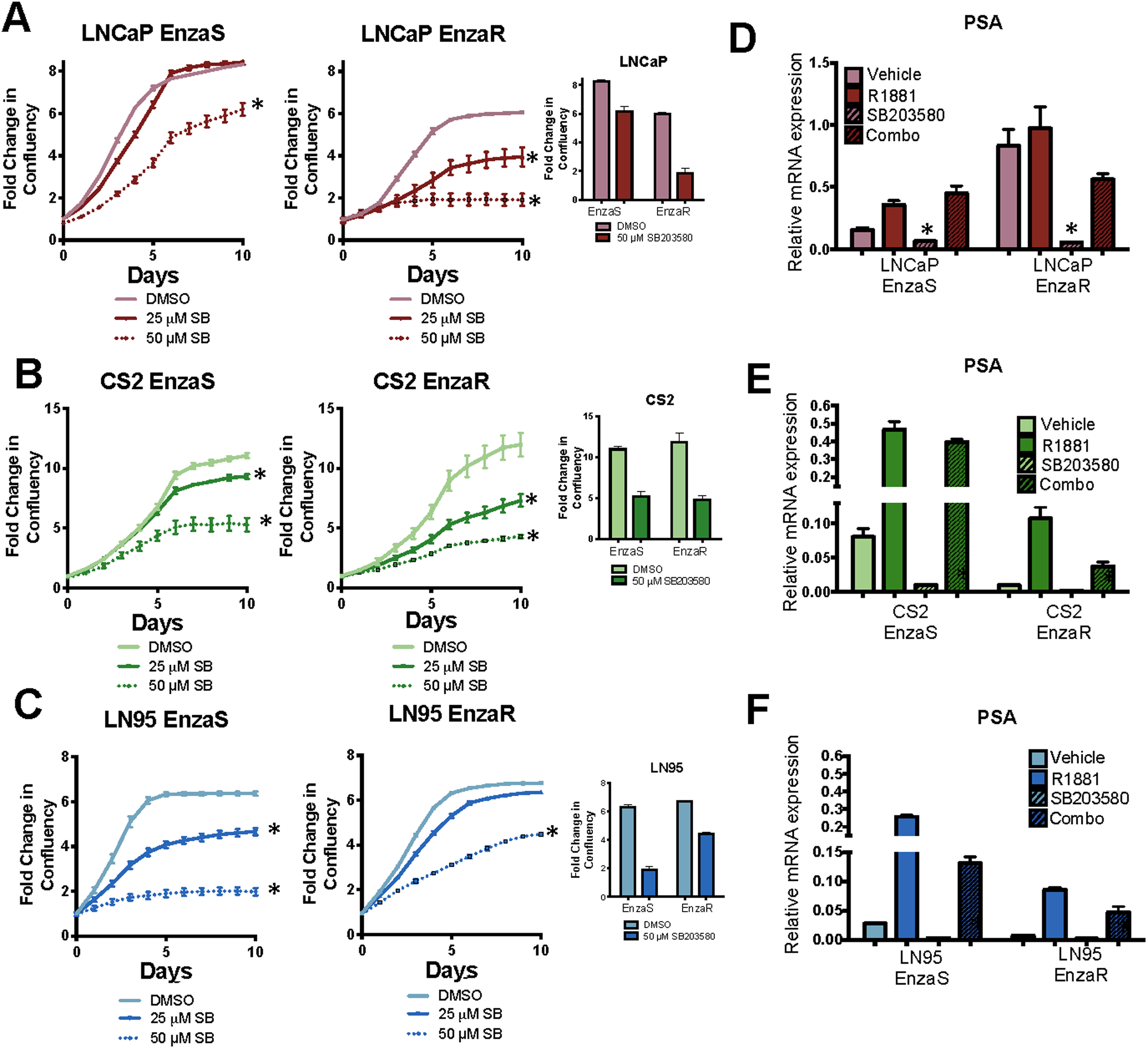
p38 inhibition is important for cell growth in EnzaR cell lines. **A-C.** Control or enzaR cells were cultured in media containing DMSO or 25μM SB203580, and cell confluence was quantified using the IncuCyte basic analyzer. **D-F.** Control or enzaR cells, treated with androgen (R1881), p38 inhibitor (SB203580), or the combination of R1881 and SB203580 were analyzed for PSA mRNA expression. Amounts were determined by qRT-PCR and normalized to GAPDH, * = p<0.05; student’s t-test.

### Activation of p38/MAPK promotes sustained AR signaling in the presence of enzalutamide

Previous studies have shown that enzalutamide induces lineage reprogramming toward an AR null, neuroendocrine-like phenotype (Ku et al., 2017; Mu et al., 2017; Paranjape et al., 2016). However, neither the RNA-Seq nor qRT-PCR analyses revealed consistent changes in neuroendocrine biomarkers across our four paired enzalutamide resistant cell lines (**Supplemental Fig. 4**). On the contrary, all enzaR cell line models maintain AR protein expression (Fig 2C, D), indicating that phenotypic shifts to AR-null (Bluemn et al., 2017) or neuroendocrine lineages (Aparicio et al., 2011; Dardenne et al., 2016) have not occurred in our four model systems. Thus, our models may recapitulate the common occurrence of AR positive metastatic castration-resistant prostate cancer post-abiraterone/enzalutamide, observed in the majority (>60-70%) of men with lethal prostate cancer (Bluemn et al., 2017) (Fig. 2). Consistent with this, treatment with androgen (R1881) induces AR activity as measured by PSA expression (Fig. 4D-F), implying that enzaR cells remain androgen dependent and responsive to AR signaling.

To understand how enzalutamide-induced p38 activation may impact AR activity, we treated control and enzaR cells with the p38 inhibitor, SB203580. Inhibition of p38 led to dramatic down regulation of AR activity in both sensitive and resistant cell lines, which suggests p38 plays a role in promoting AR activity in the setting of CRPC. Additionally, treatment with the p38 inhibitor, SB203580, partially blocked androgen-stimulated *PSA* and *NKX3.1* expression (Fig. 4D-F, **S6A**). Taken together, our data indicates p38 activation promotes convergent AR-dependent enzalutamide resistance, at least in part, by facilitating persistent AR activation.

To further validate the importance of p38 signaling in promoting enzalutamide resistant growth we performed population-level p38 knockout or p38 constitutive activation in LNCaP and LN95 cells. CRISPR/Cas9-mediated knockout of p38 delayed outgrowth of enzalutamide-resistant cells (Fig. 5A, B) and constitutive activation of p38 promoted resistance to enzalutamide treatment (Fig. 5C, D). Likewise, treatment with the p38 inhibitor, SB203560, *in vivo* significantly reduced the growth of enzaR xenografts (Fig. 4G). Consistent with the decreased tumor growth over time, histology from tumor tissues treated with the p38 inhibitor indicated acute inflammation and necrosis with little viable tumor compared to diluent (control) treated tumors (Fig 4H). Together, these data suggest a causal mechanistic relationship between p38 activation and acquired resistance to AR inhibition in prostate cancer that can be overcome by p38 blockade *in vivo* and

**Figure 5.**
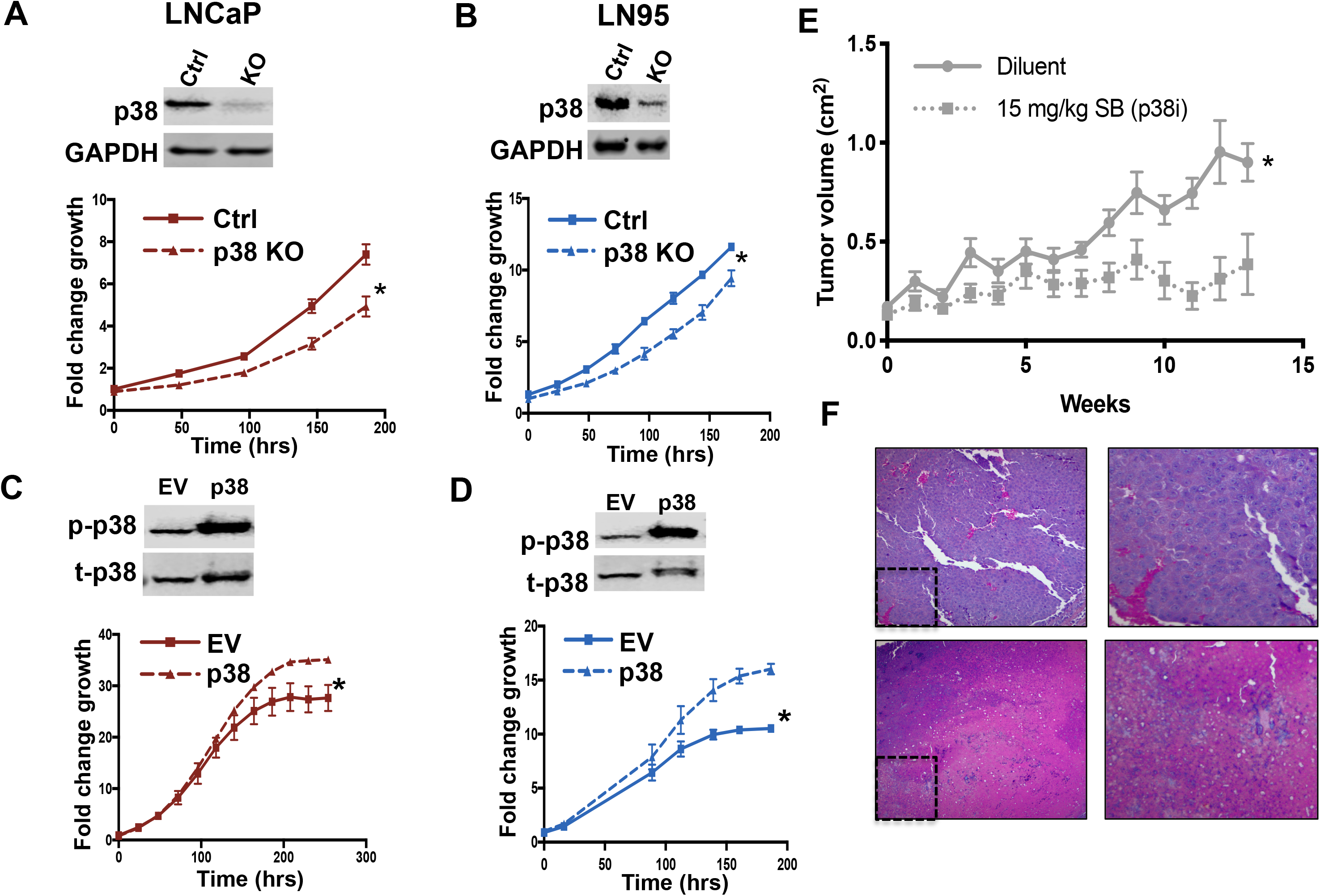
Molecular alteration of p38 expression regulates enzalutamide sensitivity. **A.** EnzaR cells with p38 knockout were cultured and cell confluence was quantified using the IncuCyte basic analyzer. Knockout of p38 in enzaR cell line populations decreases cell proliferation of LNCaP (A) and LN95 cells (B). **C.D.** Enzalutamide sensitive cells engineered to overexpress p38 were cultured and cell confluence was quantified as above. Overexpression of p38α in enzalutamide sensitive control cell lines increases cell growth in response to enzalutamide treatment. **E.** Mice were injected subcutaneously with 8 x 10^6^ CS2-enzaR cells and treated with diluent or 15 mg/kg of p38 inhibitor (SB203580). Tumor growth was measured weakly over 13 weeks. * = p<0.05; student’s t-test. **F.** Representative 10X images of H&E staining from xenograft tumors collected after treatment with diluent or p38 inhibitor (15 mg/kg SB203580) demonstrating viable (top) and necrotic (bottom) tumor.

### A p38/MAPK axis induces pro-metastatic and immuno-evasive phenotypes

The observation that p38 activity was increased in metastatic tissues suggested that p38 may facilitate metastatic progression or is upregulated during the metastatic cascade. To better understand the mechanisms by which p38 activity may promote metastasis, we interrogated known drivers of metastatic prostate cancer. One of these drivers of metastasis is Snail, a master regulator of epithelial plasticity that promotes migration and invasion and is strongly expressed in 100% of metastatic prostate cancer biopsies (Ware et al., 2016). We have previously shown that Snail activates AR nuclear localization to drive plasticity, invasion, and enzalutamide resistance (Ware et al., 2016). Snail has also been implicated in epithelial plasticity and loss of AR activity in prostate cancer models of AR therapy resistance as well through direct binding to the AR gene locus (Miao et al., 2017). We observed Snail upregulation in the p38-activated, enzalutamide-resistant cell lines (Fig. 6A) as compared to enzalutamide sensitive models, suggesting that enzalutamide-induced p38 activation induces Snail upregulation. Consistent with Snail upregulation, enzalutamide-resistant cells displayed an increase in repressive phosphorylation at serine 9 on the upstream Snail destabilizing protein and known p38 target, GSK3β (Bikkavilli et al., 2008) (**Supplemental Fig. S8A**). These results provide a mechanistic link connecting p38 activation to increased Snail through suppression of GSK3β activity, which may lead directly to altered AR activity and nuclear localization.

**Figure 6.**
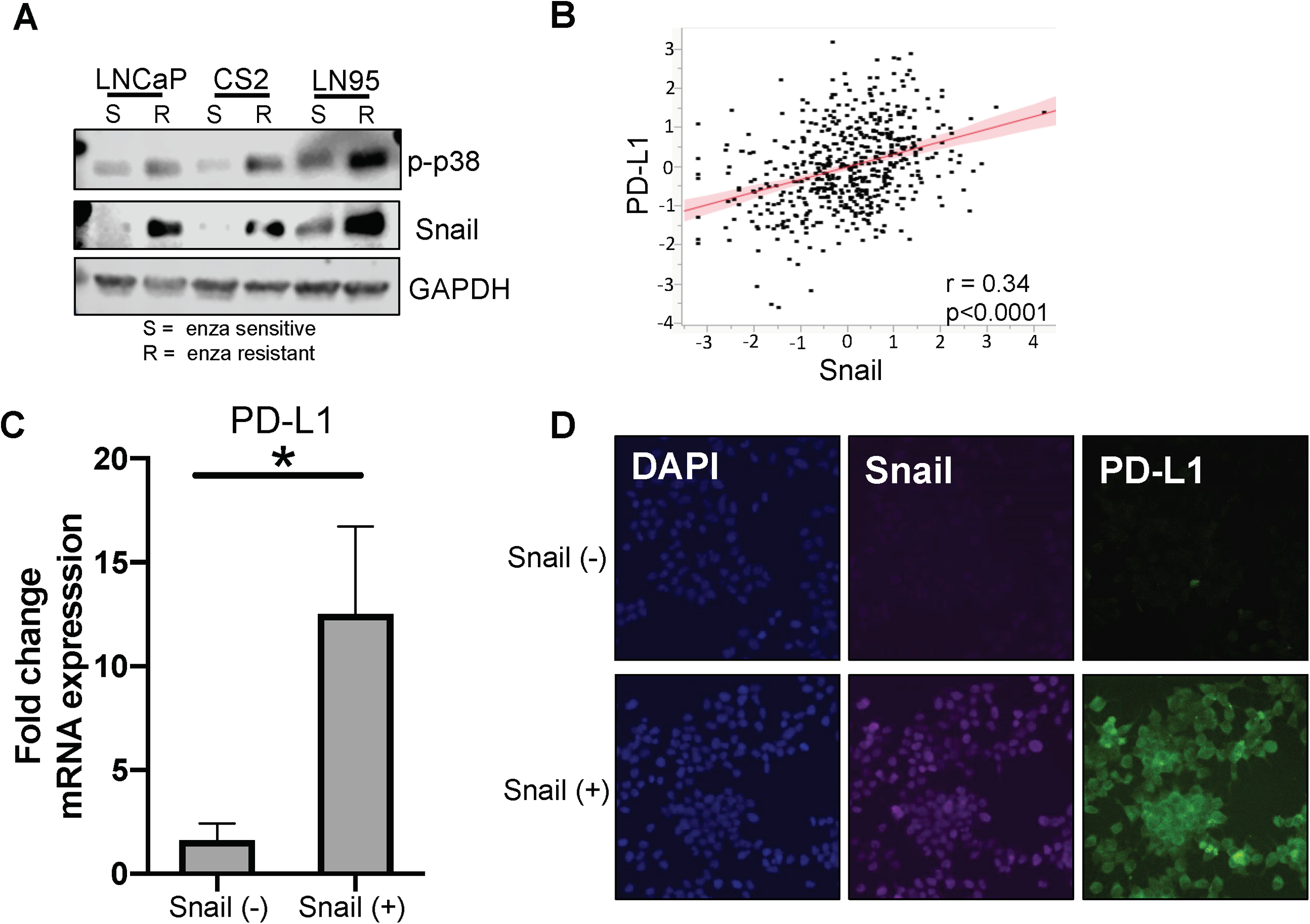
PD-L1 expression correlates with Snail expression and activity during enzalutamide resistance. **A.** Western blot analysis of phospho-p38 and Snail expression in control and enzaR cell lines. GAPDH is used as a loading control. **B.** Analysis of RNA-Seq data from The Cancer Genome Atlas reveals a significant positive correlation between Snail and PD-L1 expression in prostate cancer tissues. **C.** PD-L1 mRNA expression in LN95 cells with and without Snail activation. **D.** Snail and PD-L1 protein expression in LN95 cells with and without Snail activation. Blue: Hoechst stained nuclei; Red stained Snail; Green stained PD-L1.

Prior studies have demonstrated an association between enzalutamide resistance and PD-L1 expression on tumor and immune cells, as well as the potential benefits of combined enzalutamide and PD-1 inhibition (Bishop et al., 2015; Graff et al., 2016; Zhang et al., 2018), suggesting a connection between AR inhibition and immune evasion. Interestingly, we also observed a significant positive correlation in gene expression data in prostate cancer tissues from The Cancer Genome Atlas between Snail expression and expression of the immune checkpoint molecule, PD-L1 (Fig. 6B). To identify Snail as an effector molecule upstream of PD-L1 expression, we used an inducible system to activate Snail in LN95 prostate cancer cells (Ware et al., 2016). Snail activation led to upregulation of PD-L1 mRNA and protein (Fig. 6C, D). We also noted common upregulation of PD-L1 by reverse phase protein array analysis, qPCR, and western blotting in all enzaR models as compared to the parental models (Fig. 7A-C).

**Figure 7.**
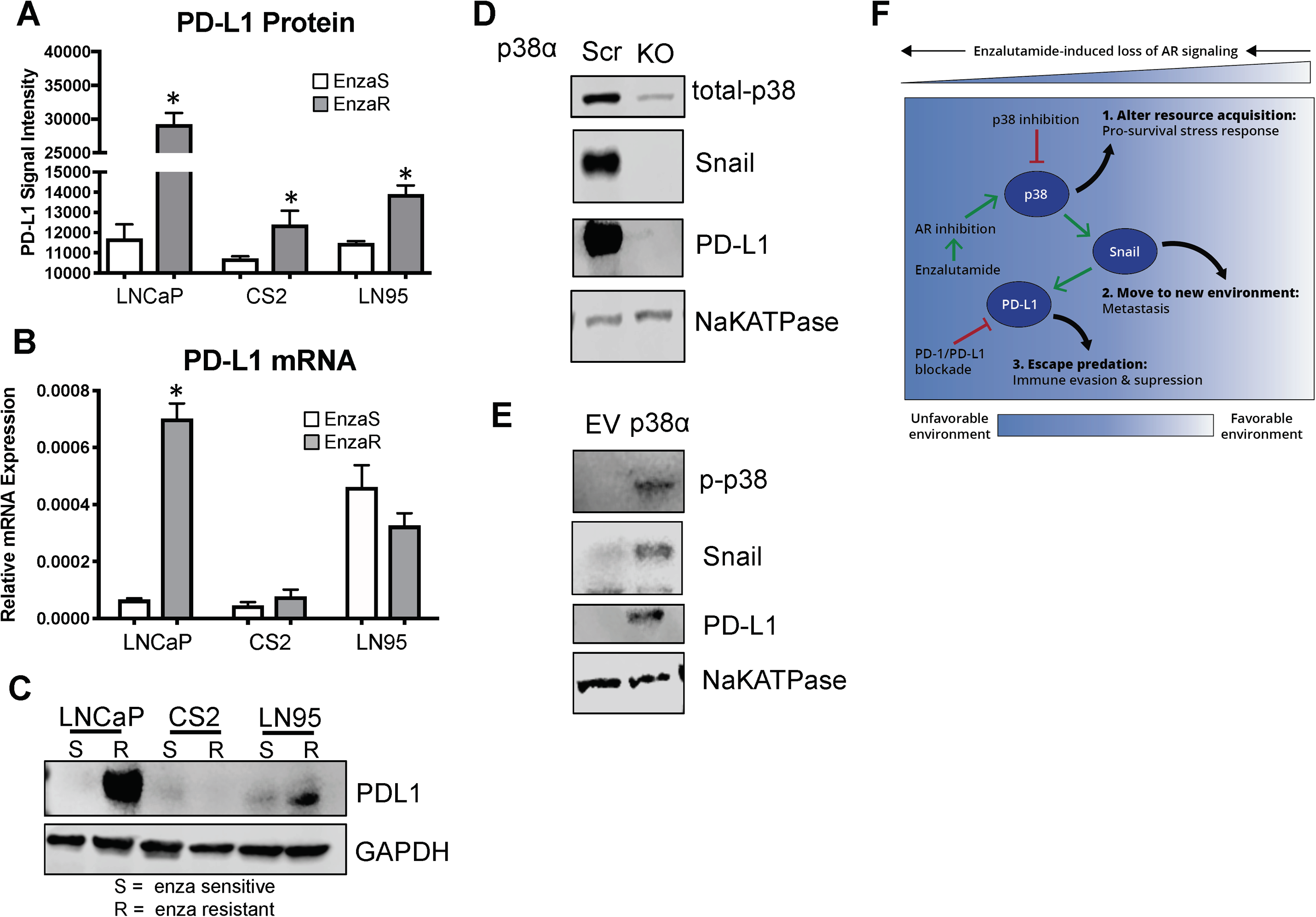
p38 is a druggable target upstream of Snail and PD-L1. **A.** PD-L1 protein expression is significantly increased in enzaR cells by reverse phase protein array analysis. **B-C.** PD-L1 mRNA (**B**) and protein (**C**) expression measured by qPCR and western blot, respectively, in enzaR cells. **D.** Western blot analysis of p38, Snail, and PD-L1 in LN95 enzaR cells with p38α knockout. NaKATPase is used as a loading control. **E.** Western blot analysis of p38, Snail, and PD-L1 in LN95 control cells with p38α activation. NaKATPase is used as a loading control. **F.** Model of p38 as a druggable target upstream of Snail and PD-L1 induced during resistance to enzalutamide.

Given the positive relationship between p38 phosphorylation, PD-L1, and Snail expression in both our models and clinical samples, we next sought to determine the mechanistic links between p38, Snail, and PD-L1. To do this we assessed the consequence of inhibiting or activating p38 signaling on Snail and PD-L1 expression. Remarkably, p38α knockdown led to downregulation of both Snail and PD-L1 (Fig. 7D) and constitutive activation of p38α induced Snail and PD-L1 upregulation (Fig. 7E). These experiments suggest that p38 activation can promote a pro-metastatic, immune evasive phenotype by stabilizing Snail expression (Ryu et al., 2019) and leading to downstream PD-L1 upregulation. Together, our data suggest that p38 may provide a common mechanistic explanation for evolutionary convergence of cancer cell survival, metastasis, and immune evasion phenotypes in AR positive metastatic castration-resistant prostate cancer, which could be therapeutically targeted through p38 blockade to treat men with hormone therapy resistant prostate cancer (Fig. 7F).

## Discussion

### Hormone therapy-resistant prostate cancer cells undergo convergent evolution onto a pro-survival, metastatic, immuno-evasive gene regulatory network

Our integrated, multi-omics analysis of the evolution of enzalutamide resistance in prostate cancer revealed consistent activation of the p38/Snail/PD-L1 gene regulatory network in the context of persistent AR expression and activity. Activation of this gene expression pathway was observed in the face of heterogeneous and unique acquired genetic landscapes across the diverse model systems tested. Likewise, our current and previous clinical analyses revealed upregulation of both p38 and Snail in metastases from metastatic castration-resistant prostate patients (Ware et al., 2016), all of which are likely to harbor substantial spatial and temporal genetic heterogeneity. This phenotypic similarity is reminiscent of convergent evolution in natural systems in which strong selective pressures lead to optimal adaptive phenotypes, such as the independent evolution of flight and the loss of sight in low-light environments across multiple, genetically-distinct taxonomic lineages (Gatenby et al., 2011). These results suggest that, in addition to known genetic drivers of resistance (Azad et al., 2015; Ku et al., 2017; Mazrooei et al., 2019; Mu et al., 2017; Romanel et al., 2015), the evolution of resistance can also be mediated by gene regulatory phenotypes rather than any common genetic driver. This convergent evolution into critical molecular driver pathways may be overlooked through the analysis of DNA or RNA-based sequencing approaches in isolation.

### The p38/Snail signaling axis promotes cell-intrinsic enzalutamide resistance and metastasis

Eukaryotic cells have evolved to respond to environmental stressors through a robust and highly-coordinated stress response system (Sun and Zhou, 2018). This system is dynamically regulated through multi-level regulation of gene expression networks, which induce reversible phenotypic changes to enable cellular responses.

In the context of cancer, cells encounter a range of environmental stressors, such as hypoxia, reactive oxygen species, glucose and nutrient deprivation, and mechanical stress (reviewed in (Chen and Xie, 2018). In addition to these microenvironmental factors, cancer therapy dramatically alters the environment and induces a range of cellular responses to enable cell survival in the face of therapy-induced resource depletion. One of these key cellular responses is p38/MAPK pathway activation. The p38 pathway is an evolutionarily-conserved stress response system (Li et al., 2011a), which is responsible for transducing stress signals, such as DNA damage, reactive oxygen species, cytokines, and changes in osmotic pressure, from the extracellular environment to activate transcriptional response programs (Coulthard et al., 2009). Activation of the p38/MAPK pathway has been implicated in numerous cancers, including metastatic prostate cancer (Drake et al., 2016; Khandrika et al., 2009; Koul et al., 2013). Consistent with its role in mediating cellular stress responses, we found that p38 promotes cell-intrinsic resistance to enzalutamide by promoting reactivation of AR activity. Mechanistically, we observed increased phosphorylation of GSK3β at serine 9 (**Supplemental Fig. 8**), which leads to inactivation of GSK3β activity and induces upregulation of Snail. This model is supported by previous reports, which have demonstrated that p38 inhibits GSK3β through phosphorylation (Thornton et al., 2008), which can lead to Snail stabilization (Zhou et al., 2004). Importantly, we have demonstrated that Snail contributes to enzalutamide resistance via upregulation of AR activity (Ware et al., 2016). In addition to its functions in enzalutamide resistance, p38 and Snail also drive a pro-metastatic and death-resistant phenotype in prostate cancer. Consistent with this, inhibition of p38 leads to reduced survival, clonogenicity, and invasion in prostate cancer cells (Khandrika et al., 2009). Others have also shown that activation of p38 induces TGF-β-mediated Snail upregulation and invasion (Medici et al., 2011). Similarly, we previously demonstrated that Snail is strongly elevated in metastatic, castration-resistant prostate cancer compared to primary prostate cancer (Ware et al., 2016). We also found that Snail activation leads to enhanced migration and invasion of prostate cancer cells (Ware et al., 2016). In the current study, we define a mechanistic connection between the p38 pathway and Snail that drives enzalutamide resistance and metastasis.

### The p38/Snail axis promotes cell-extrinsic immune evasion

Coupled with their roles in enzalutamide resistance, our data suggest the p38/Snail pathway may also contribute to immune evasion by upregulating the immune checkpoint molecule, PD-L1. In addition, enzalutamide exposure alone leads to upregulation of Snail and PD-L1 (Bishop et al., 2015), and our data demonstrates Snail activation can upregulate PD-L1 expression. Consistent with this, PD-L1 is upregulated on melanoma cells through activation of p38 (Noh et al., 2015), providing further strong support for our hypothesis that the p38/Snail axis drives PD-L1 expression. Similarly, in both chronic viral infections and cancer, p38 activation has been linked to inhibition of Stimulator of IFN genes (STING) expression, enhanced CXCR2-mediated myeloid derived suppressor cell activity, and immune evasion (Chen et al., 2017; Wang et al., 2006; Zhang et al., 2017). Together, these findings suggest that p38 and/or immune checkpoint blockade may be therapeutically efficacious in the post-enzalutamide setting. In support of this notion, Graff et al. observed rapid reductions in prostate specific antigen and radiographic responses in approximately ~20% of men with enzalutamide resistant metastatic castration-resistant prostate treated with a combination of the anti-PD-1 therapy pembrolizumab and enzalutamide (Graff et al., 2016; Graff et al., 2018). Furthermore, PD-L1 can indirectly activate p38 through DNA-PKcs, which promotes chemoresistance (Wu et al., 2018), and therefore suggests a common feed forward loop that may connect PD-L1 expression, DNA repair and pro-metastatic pathways (Goodwin et al., 2015), and convergent p38 activity. Interestingly, PD-L1 expression in the tumor and tumor microenvironment is also associated with poor outcomes and an aggressive and metastatic phenotype in many cancer subtypes (Kim et al., 2016; Wang et al., 2018).

### Beyond enzalutamide resistance

The current work highlights the role of the p38 pathway in promoting enzalutamide resistance in prostate cancer. However, the importance of the p38 pathway as a general stress response to any unfavorable environment may suggest the potential of targeting the p38 pathway in other resource-limiting settings in cancer. Indeed, our clinical data suggest that the p38/Snail pathway is induced during metastasis even prior to androgen deprivation or enzalutamide treatment. Therefore, tumor cells undergoing a stress response independent of enzalutamide or prostate cancer may also rely on p38 activation to persist and survive in ecologically-unfavorable environments, such as the hostile environment of the bloodstream during dissemination, or during treatments that target other oncogenic drivers across different cancer types. For example, p38 activity has been shown in both breast and lung cancer to promote chemotherapy resistance (Flem-Karlsen et al., 2019; Liu et al., 2016; Lu et al., 2018). Activation of p38 has also been linked to DNA repair (Canovas et al., 2018), which is also enriched in our models of enzalutamide resistance (Fig. 3B). Therefore, p38 inhibition in combination with chemotherapeutic drugs that induce chromosome instability may have therapeutic potential. These data suggest there may be clinical utility in re-purposing p38 inhibitors for the investigation of reversing treatment resistance in metastatic castration-resistant prostate and other cancers (Fig. 7F).

## Acknowledgements

Funding: This work was supported by grants NIH F32 CA192630 (K.W. Ware), NIH 1R01CA233585-01 (A.J. Armstrong), NIH P30 CA014236 (M. Kastan), Department of Defense W81XWH-18-1-0189 (J.A. Somarelli), Triangle Center for Evolutionary Medicine (J.A. Somarelli), Astellas Scientific and Medical Affairs (A.J. Armstrong), and the Prostate Cancer Foundation (A.J. Armstrong; J.A. Somarelli). Prostate Cancer Foundation Young Investigator Award (J.M. Drake) The authors thank Drs. William Eward and Everardo Macias for sharing their IncuCyte systems to monitor cell growth and apoptosis over time as well as the core research facilities involved in this work (Duke Functional Genomics, Duke Biorepository & Precision Pathology, Duke Genomic Analysis and Bioinformatics). We would also like to thank Medivation/Astellas for providing the AR inhibitor, enzalutamide.

## Author Contributions

Conceptualization, KEW, JAS, AJA; Methodology, KEW, JAS, AJA, DLC, MP, EFP, ZS, JMD, LAC, WCF; Investigation, KEW, JAS, SG, JE, GK, BJP, RGA, DR, BCT, AA, MP; Writing-Original Draft, KEW, JAS, AJA; Visualization, MUS, JAS, KEW; Resources, JAS, AJA, KEW, JF, SRP, TZ, SG, JMD

## Materials and Methods

### Patient samples

For metastatic prostate cancer samples (n=20) and a second cohort of localized prostate cancer samples (n=30), formalin fixed metastatic tissue was collected from the Duke University pathology department and Duke Cancer Institute Biorepository Core under a separate Duke IRB approved protocol. Clinical data on prior therapy and metastatic site were collected.

### Cell lines

MDA-PCa-2b and LNCaP cells were obtained from ATCC using the Duke University Cell Culture Facility. MDA-PCa-2b cells were cultured in F12-K media supplemented with 20% fetal bovine serum (Sigma), 1% penicillin/streptomycin streptomycin (Life Technologies), cholera toxin, epidermal growth factor, phosphoethanolamine, hydrocortisone, sodium selenite, and insulin as recommended by ATCC. LNCaP cells were cultured in RPMI containing 10% fetal bovine serum and 1% penicillin/streptomycin. LNCaP95 cells were kindly provided by Dr. Scott Dehm (University of Minnesota) and are reported in previous studies (Liu et al., 2013; Ware et al., 2016). LNCaP95 and CS2 cells are androgen-independent cell lines derived from the parental LNCaP cells. LNCaP95 and CS2 were cultured in RPMI containing 10% charcoal stripped fetal bovine serum and 1% penicillin/streptomycin. Resistant cell populations were cultured with the addition of 50 μM enzalutamide (provided by Medivation/Astellas). Enzalutamide resistant cells (EnzaR cells) were generated by chronic culture with increasing doses of enzalutamide to a concentration of 50 μM. Cells were authenticated and re-authenticated following enzalutamide resistant through sequencing including the presence of known LNCaP AR ligand binding domain mutation T877A. Cells stably expressing inducible Snail (Addgene plasmid #18798), constitutively active p38 or MAPK14 gRNA targets were generated by transduction of cells as described previously (Mani et al., 2008).

### Cell growth and viability assays

Control and Enza-R cells were cultured in media containing DMSO (Sigma) or 50 μM enzalutamide for at least one week. Cells were counted using the Countess II (Life Technologies), 2500 cells were seeded in a 96-well plate. Cell confluence was monitored using the IncuCyte live cell analysis system (Essen Biosciences) and standard error of the mean (SEM) was calculated from triplicate wells. For cells stably expressing p38 or MAPK14 gRNA, cells were treated with DMSO or 25 μM of SB203580 (selleckchem) and monitored as above. For cell viability, apoptosis was measured after 10 days of drug treatment using the IncuCyte Caspase-3/7 green apoptosis assay reagent and quantified using the IncuCyte basic analyzer.

### RNA-seq

Total RNA was isolated using the Quick-RNA Miniprep kit (Zymo Research). RNA-seq, total RNA was submitted to the Duke Center for Genomic and Computational Biology core for sample preparation, sequencing, and analysis. RNA-seq data was processed using the TrimGalore toolkit (http://www.bioinformatics.babraham.ac.uk/projects/trim_galore) which employs Cutadapt^1^ to trim low quality bases and Illumina sequencing adapters from the 3’ end of the reads. Only pairs where both reads were 20nt or longer were kept for further analysis. Reads were mapped to the GRCh37v73 version of the human genome and transcriptome^2^ using the STAR RNA-seq alignment tool^3^. Reads were kept for subsequent analysis if they mapped to a single genomic location. Gene counts were compiled using the HTSeq tool (http://www-huber.embl.de/users/anders/HTSeq/). Only genes that had at least 40 reads in any given library were used in subsequent analysis. Normalization and differential expression was carried out using the EdgeR^4^ Bioconductor^5^ package with the R statistical programming environment (www.r-project.org). A negative binomial generalized log-linear model^6^ was used to identify differentially expressed genes between the different genotypes when comparing against specific control samples. Enriched pathways were determined by GSEA^7^ for each comparison.

### Real-Time quantitative RT-PCR

For qPCR, total RNA was reverse transcribed using the High-Capacity cDNA Reverse Transcription Kit (Life Technologies). Aliquots of 5-fold diluted reverse transcription reactions were subjected to quantitative (q)PCR with KAPA SYBR FAST master mix using the Vii7 real time-PCR detection system (Applied Biosystems). GAPDH mRNA levels were measured for normalization, and the data are presented as “Relative Expression”. A complete list of primer sequences is provided in the supplementary text.

### TCGA Analysis

The results published here are in part based upon data generated by The Cancer Genome Atlas (TCGA) Research Network: http://cancergenome.nih.gov. Data available from TCGA was analyzed using the Kruskal Wallis test to evaluate the correlation between Snail expression and PD-L1 expression from prostate cancer patients.

### Reverse phase protein array

Cells were seeded (300,000/well) in 6-well plates and allowed to incubate for 5 days. Cells were then rinsed with PBS, flash frozen, and analyzed as previously described (Baldelli et al., 2017; Pierobon et al., 2017).

### Immunoblot analyses

For immunoblot analysis cells extracts were mixed with SDS sample buffer and submitted to SDS-PAGE. Following electrophoretic transfer onto nitrocellulose, the filters were blocked in Starting Block (Thermo), incubated with antibodies and developed using the Odyssey-FC imager (LI-COR).

A complete list of primary antibodies and their dilutions is provided in the supplementary text.

### Immunofluorescence staining

For cells expressing inducible Snail, cells were pretreated with ethanol (EtOH) or 4OHT. For immunofluorescence (IF) staining, cells were fixed in 4% PFA, permeabilized with 0.2% Triton X-100, and stained with Hoechst. Cells were blocked with 5% bovine serum albumin (BSA, Sigma) prior to incubation with primary antibodies. Cells were incubated in Alexa Fluor secondary antibodies (Life Technologies) and then imaged on an inverted Olympus IX 73 epifluorescence microscope.

### Immunohistochemistry

We performed antibody optimization and analytic validation for all antibodies as previously described (Armstrong et al., 2016). An expert prostate cancer pathologist blinded to outcomes evaluated antibody staining in parallel with hematoxylin and eosin. Scoring of each biomarker used a 0 to 3 scale for both intensity and a <25%, <50%, <75%, <100% scale for frequency of expression in each tumor sample.

### Animal Experiments

Six to eight week male mice were injected subcutaneously with 8 x 10^6^ CS2-enzaR cells. Prior to injection cells were resuspended in 50% matrigel supplemented with 10%FBS/RPMI. Mice were treated with diluent or 15 mg/kg of p38 inhibitor (SB203580). Tumor growth was measured using calipers weakly over 13 weeks.

### Statistical Analysis

Data are shown as means ± SEM. Student’s t-test or multiple group comparison was performed by one-way ANOVA followed by the Sidak method for comparison of means. P≤0.05 is considered significant. Differences in phospho-p38 expression between localized and metastatic samples were analyzed using a Chi-square test. Data available through the TCGA was analyzed using the Kruskal Wallis test using JMP (version pro 12). All other analyses were performed using Prism (version 8.0d).

## Supplemental Information Titles and Legends

**Figure 1**

**Treatment with enzalutamide enriches for the F876L agonist mutation in LNCaP enzaR cells.** Alignment of RNA-seq tracks using the integrative genomic viewer.

**Figure 2**

**Copy number alterations in control versus enzaR cell lines.** Array comparative genomic hybridization from genomic DNA isolated from paired enzaR cell line models. Alteration losses and gains greater than one log are highlighted in blue or red, respectively.

**Figure 3**

**EnzaR cell lines do not undergo NEPC/Stemness transitions. A.** Representative images of enzalutamide sensitive and resistant cell lines. **B.** qPCR expression analysis of driver genes involved in neuroendocrine and stem cell pathways.

**Figure 4**

**Copy number alterations do not converge between enzaR cell lines.** Venn diagrams comparing copy number gains and losses between control and enzaR cell lines.

**Figure 5**

**EnzaR cell lines acquire very few nucleotide variants during adaptation to enzalutamide. A.** Venn diagrams comparing nucleotide variants between paired control and enzaR cell lines. **B.** Venn diagram comparing common nucleotide variants observed in enzaR compared to control cell lines.

**Figure 6**

**AR activity is inhibited by p38 inhibition. A.** Control or enzaR cells, treated with androgen (R1881), p38 inhibitor (SB203580), or the combination of R1881 and SB203580 were analyzed for NKX3.1 mRNA expression. Amounts were determined by qRT-PCR and normalized to GAPDH. **B.** Western blot analysis of phospho-p38 expression in control and enzaR cell lines with acute enzalutamide (10 μM) treatment.

**Figure 7**

**EnzaR cells display markers of a dormant phenotype. A.** Ratios of phospho-ERK to phospho-p38 protein levels. **B.** Western blot analysis of p21 expression in control and enzaR cell line models. **C.** Beta-galactosidase activity in response to enzalutamide treatment in LNCaP cells.

**Figure 8**

**Molecular alteration of p38 expression regulates enzalutamide sensitivity. A.** GSK3β serine 9 phosphorylation is significantly increased in enzaR cells by reverse phase protein array analysis.

