## Supplemental Figures for "Convergent evolution of p38/MAPK activation in hormone resistant prostate cancer mediates pro-survival, immune evasive, and metastatic phenotypes"

**Supplemental Figure 1**

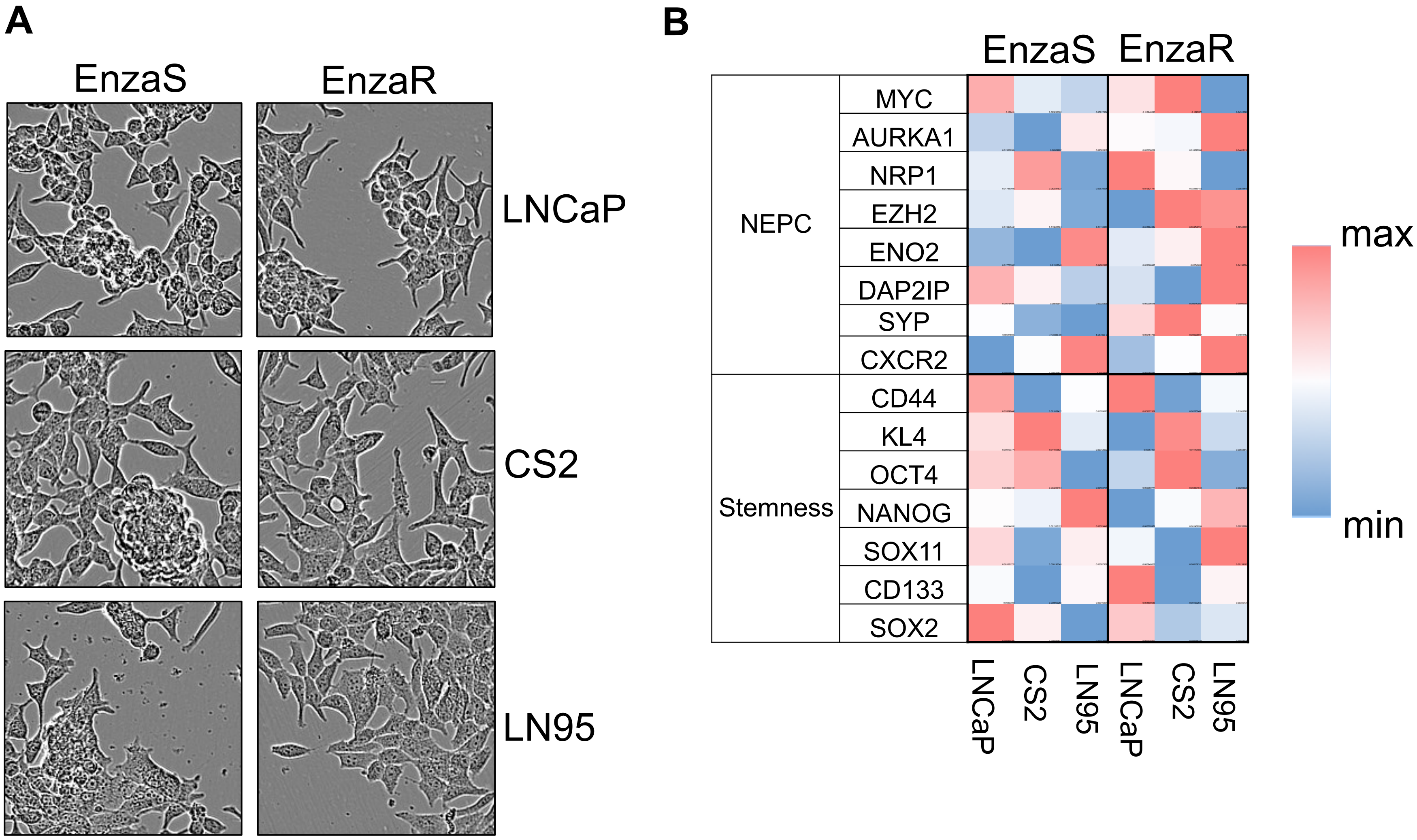

**Supplemental Figure 2**

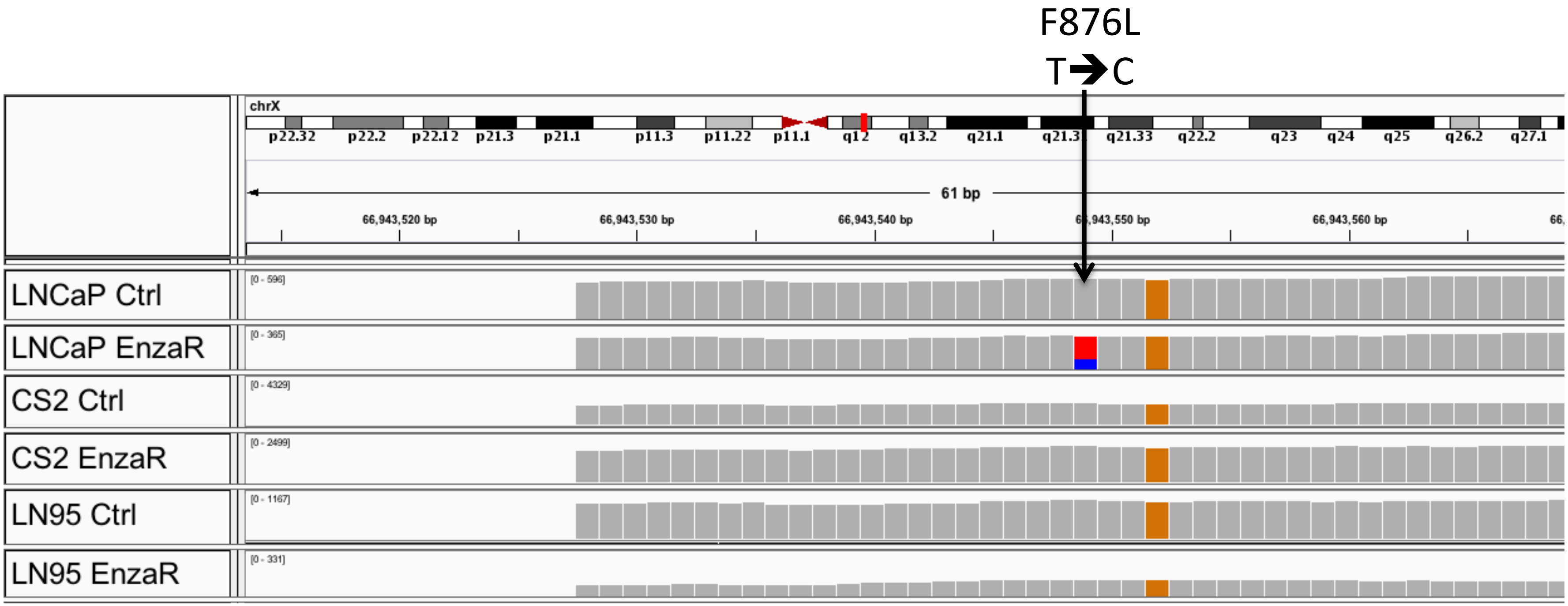

#### Supplemental Figure 3

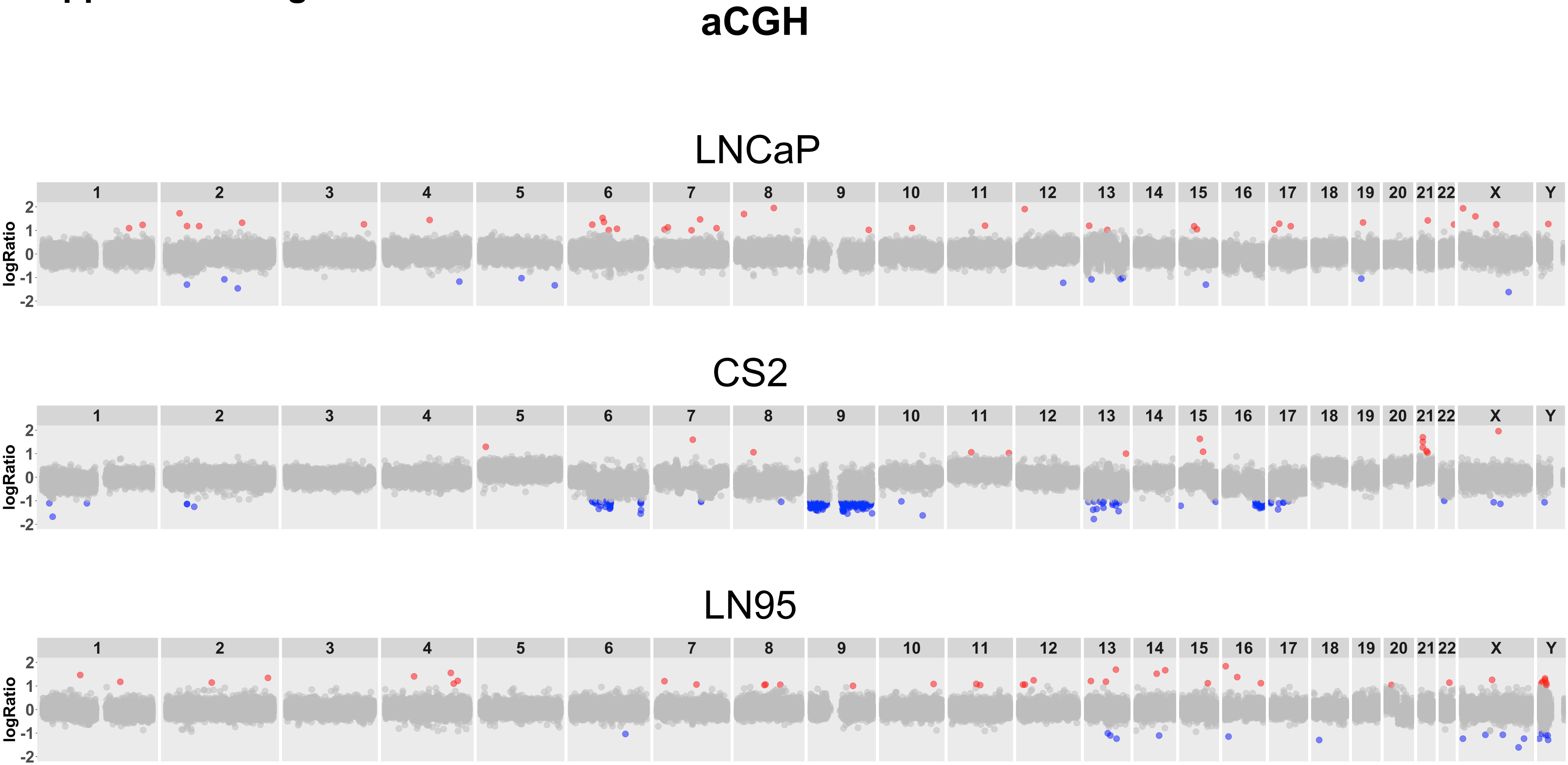

**Supplemental Figure 4**

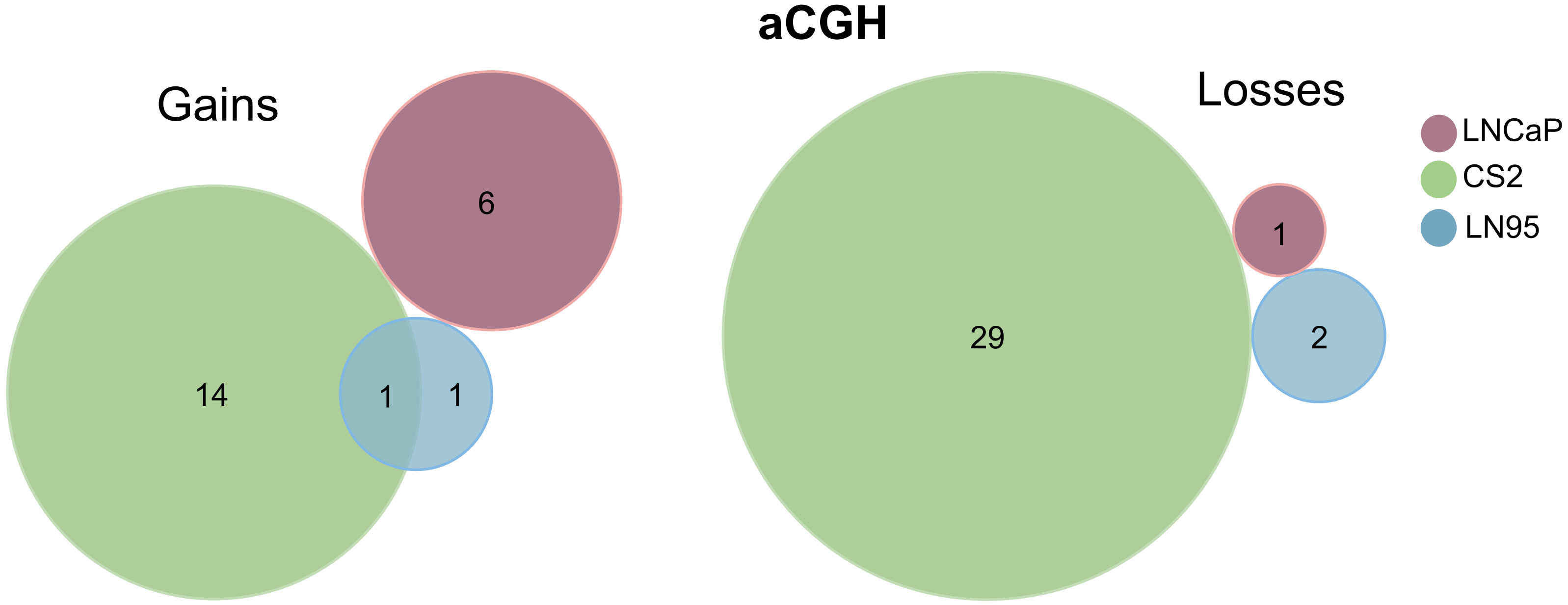

**Supplemental Figure 5**

**A**

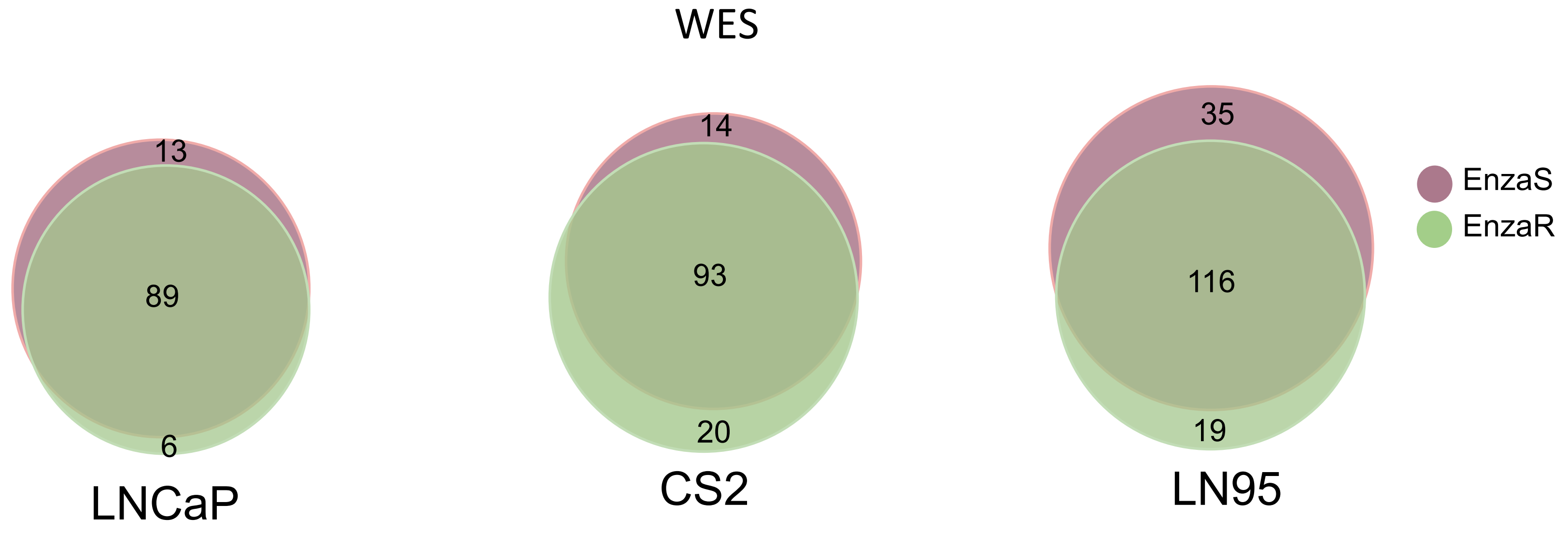


**B**

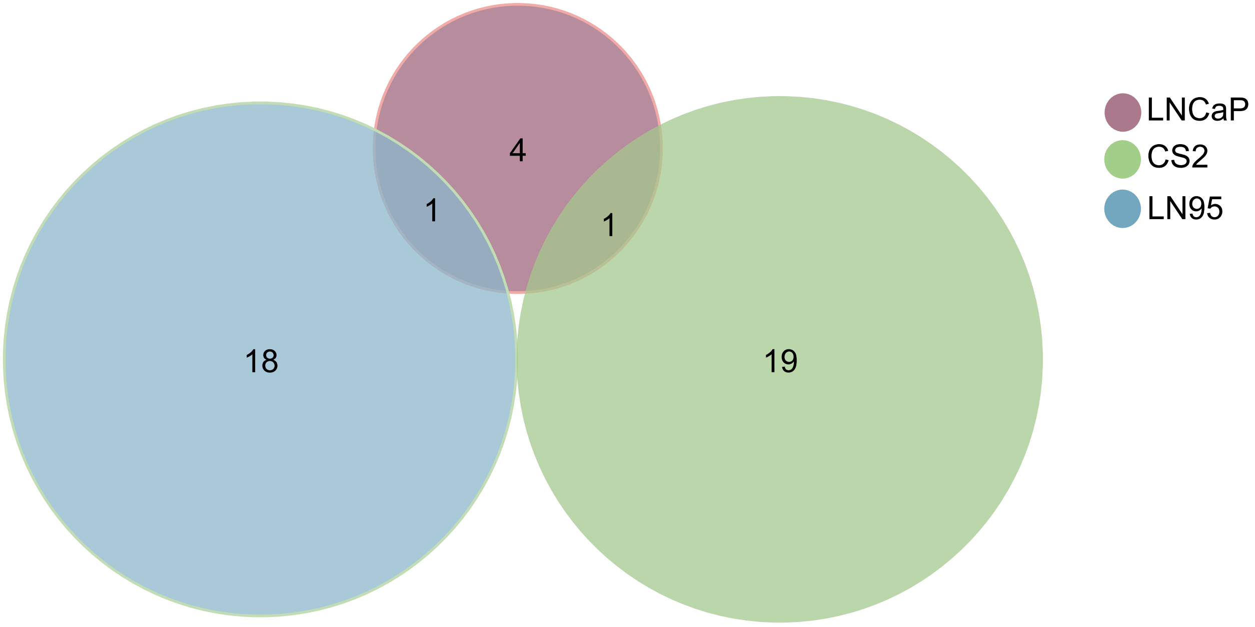

**Supplemental Figure 6**

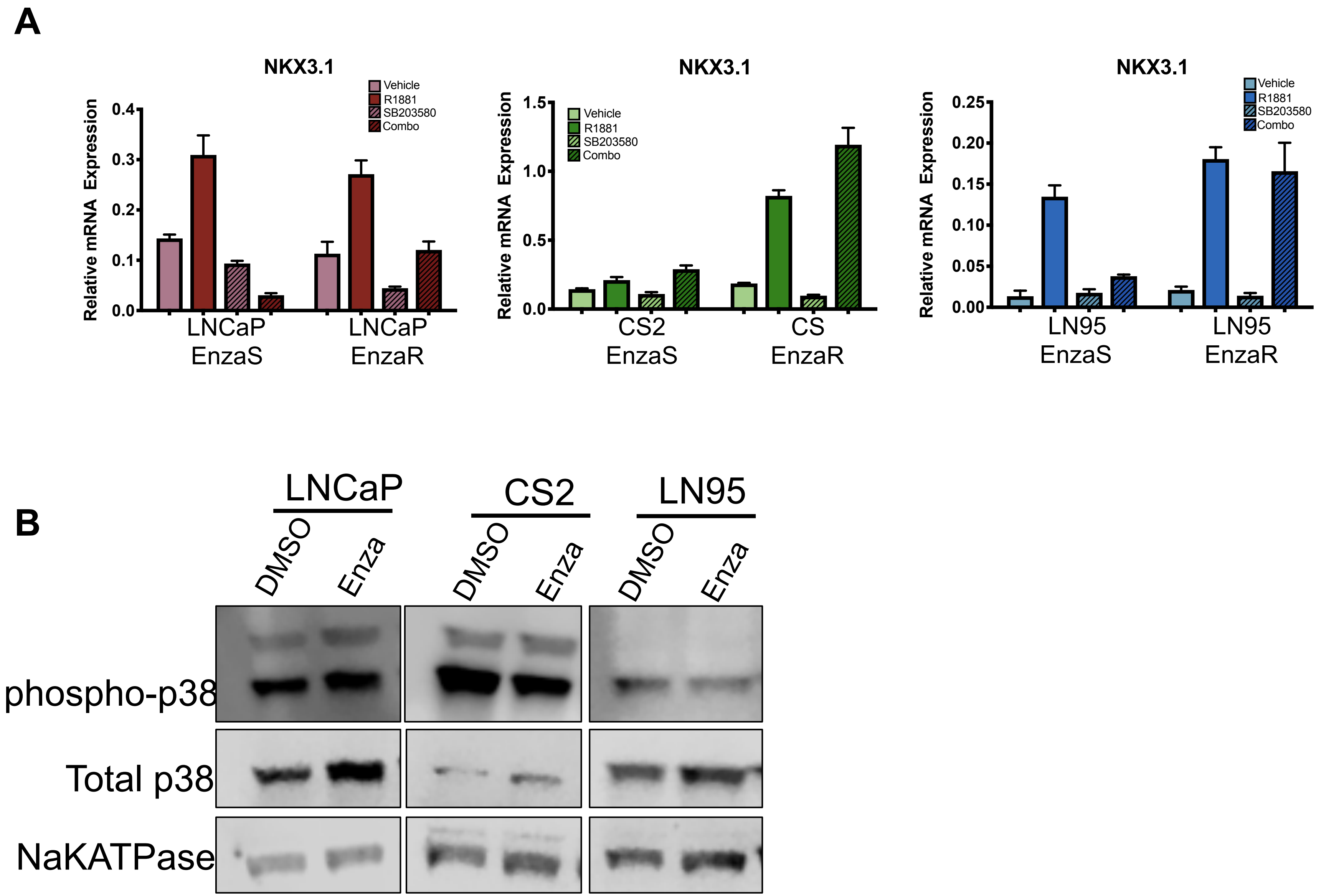

**Supplemental Figure 7**

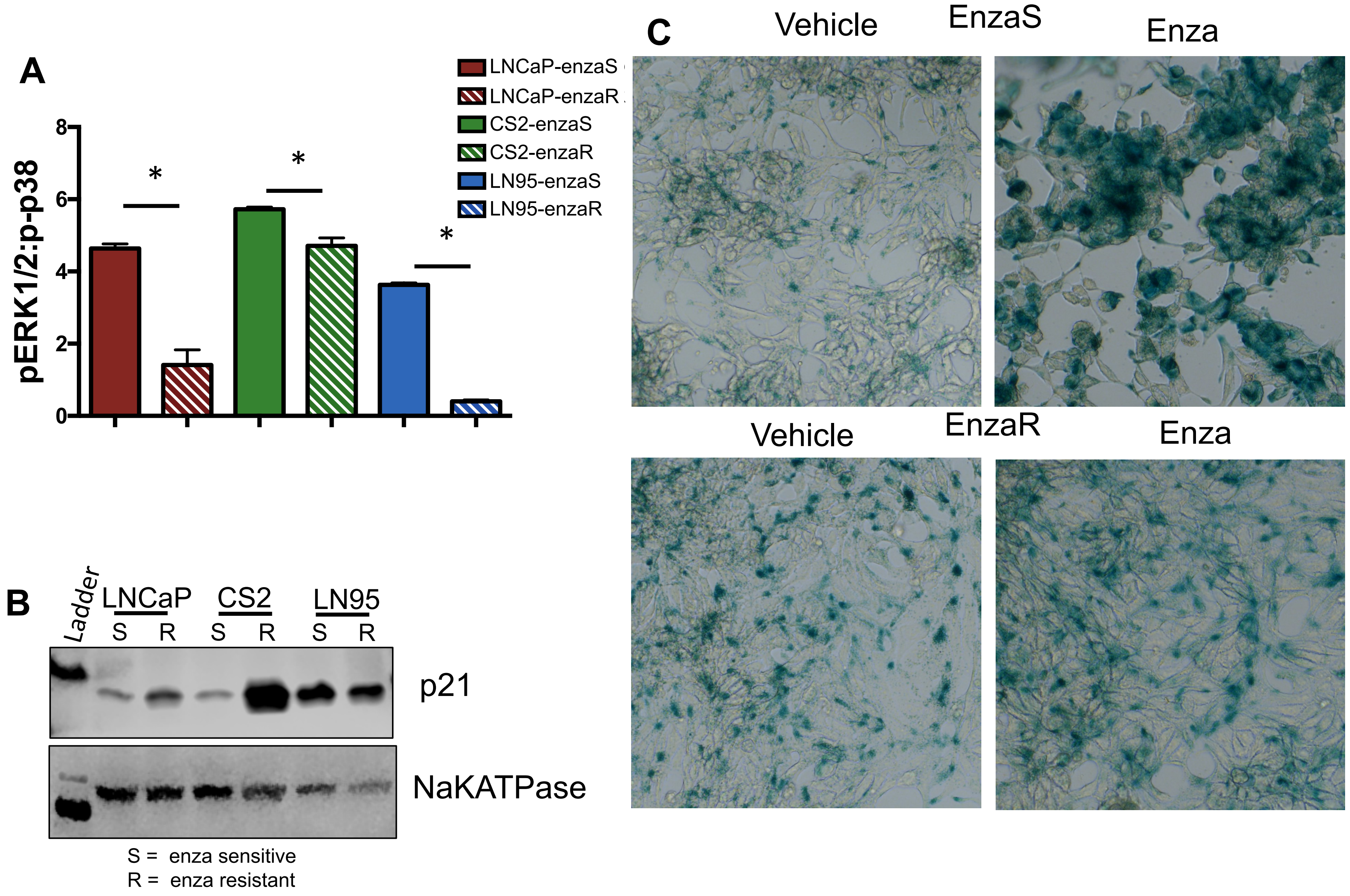

### Supplemental Figure 8

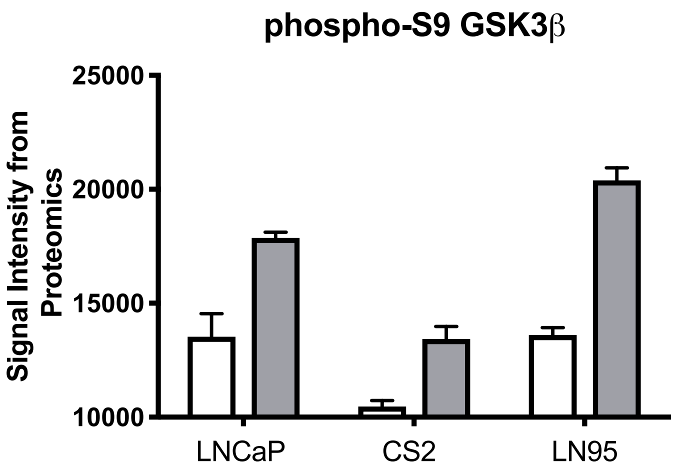
